# SSB toolkit: from molecular structure to subcellular signaling pathways

**DOI:** 10.1101/2022.11.08.515595

**Authors:** Rui Pedro Ribeiro, Jonas Goßen, Giulia Rossetti, Alejandro Giorgetti

## Abstract

We present, here, an open-source systems biology toolkit to simulate mathematical models of the signal-transduction pathways of G-protein coupled receptors (GPCRs). By merging structural macromolecular data with systems biology simulations, we developed a framework to simulate the signal-transduction kinetics induced by ligand-GPCR interactions, as well as the consequent change of concentration of signaling molecular species, as a function of time and ligand concentration. Therefore, this tool brings to the light the possibility to investigate the subcellular effects of ligand binding upon receptor activation, deepening the understanding of the relationship between the molecular level of ligand-target interactions and higher-level cellular and physiologic or pathological response mechanisms.

---

Computational systems biology approaches for modeling signaling pathways together with pharmacokinetics and pharmacodynamics mechanistic models have been making important contributions to neuroscience drug discovery and development ^1,2^. They have been used, for instance, to translate drugs’ modes of action from *in vitro* to *in vivo*, but also from animal models to human^3^. Through such models it is possible to integrate and predict important quantitative and qualitative parameters such as the drug concentration profile in blood or at the site of action and cellular signaling downstream the targets sites^4,5^. The incorporation of physiochemical-based macromolecular structure parameters in such computational systems biology approaches can not only deepen our understanding on the molecular mechanisms of drug modes of action, but also give insight in how genetic mutations can affect the subcellular downstream signaling events by altering receptor’s ligand binding, activation and signaling^4,5^.

Although the number of mathematical models of signal-transduction pathways for a variety of systems, as well as the number of tools for conducting systems biology studies and the number of computational methods to derive quantitative structure-kinetics relationships have been increasing over the years^6^, the use of systems biology simulations to study pharmacological models at the molecular level is still in its early stages. This is mostly due to the lack of a common language to annotate, exchange, reuse and update biochemical signaling pathways models, the absence of an interrelationship framework between structural and systems biology, the current limitations in reproducing kinetic parameters with molecular computational approaches and the lack of kinetic parameters needed to feed the network. Ideally, such parameters should be determined experimentally under relevant conditions to the model, but, actually the existing parameters are distributed in literature and relate to different experimental conditions^7,8^.

Here, we present the Structural System Biology (*SSB) toolkit*: an open-source systems biology object-oriented library to specifically simulate mathematical models of the signal-transduction pathways of GPCRs. The SSB toolkit allows to investigate the concentration changes of molecular species throughout signaling pathways as a function of time and ligand concentration, in order to support the comparison of signal-transduction kinetics. We envision the use of this toolkit as a scaffold that holds together ligand/structure data and systems biology simulations to study GPCRs’ modes of action.

The SSB toolkit was designed to use structure biology data to feed systems biology simulations in order to predict the effect of ligand-targets interactions on the GPCRs molecular signaling responses. Experimental or computed affinities (*K*_*d*_) or kinetic parameters (*k*_*on*_ and *k*_*off*_), can then be used to fire up the mathematical models of the signaling pathways, that predict the variation of concentration of molecular species of the pathways over time, and deduce, in the end, dose-response curves of the ligand-GPCRs interactions (Fig. 1). In this way, the SSB toolkit facilitates the comparison of the signal-transduction kinetics induced by those interactions.

**Figure 1.**
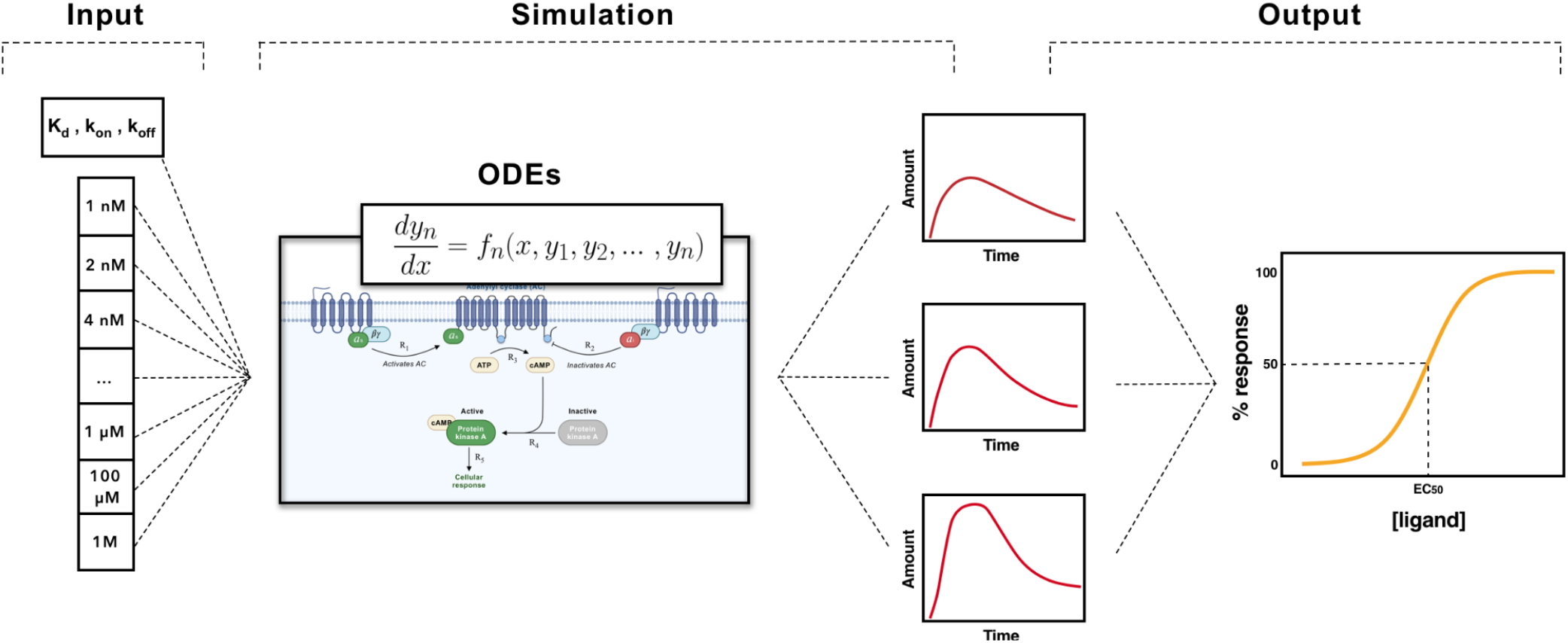
Conceptual scheme for predicting dose-response through signaling pathways’ simulation. For each concentration value the signaling model is simulated, obtaining in the end several curves of the concentration of a specific species of the pathway as a function of time. After, the maximum value of each curve is selected and plotted, resulting, in the end, in the dose-response curve.

The core of the SSB toolkit consists of the mathematical models of the main GPCRs’ signaling pathways: G_s_, G_i/o_, and G_q/11_. To implement them in our framework we adapted pre-existing models. The G_s_ and G_i/o_ pathways were implemented based on the model proposed by *Nair et al*.^9^, whereas the G_q/11_ pathway was based on the model proposed by *Chang et al*.^10^.

All the three signaling pathways were programmed using the PySB^11^ python library, designed specifically for running systems biology simulations. In detail, we start by defining all the species, initial concentrations, reactions, and reactions constants. Then, all the ordinary differential equations (ODEs) are integrated and the variation of the concentration of the molecular species that make part of the model can be monitored throughout the time evolution of the ODEs. To obtain a dose-response curve individual simulations of the pathway according to a range of ligand concentrations must be performed (Fig. 1). The dose-response curve is, then, obtained by fitting a logistic regression to the maximum concentration value from each individual simulation. In the end, a curve of the response in function of the ligand concentration is obtained (Fig. 1), from which the ***efficacy*** parameter of a ligand can be extracted.

Another novel feature of the SSB toolkit is the possibility to investigate the parameters of the model that leads to a better fit of the model to experimental data.

The open-source SSB toolkit was implemented as an object-oriented python module, which grants its readability, reusability, and deployment. It was developed having in mind its use through *Jupyter* notebooks, nowadays broadly adopted by the scientific community. In this way, users are allowed to create workflows according to their needs.

To start a SSB simulation, experimental or computed *K*_*d*_, *k*_*on*_ or *k*_*off*_ parameters must be provided as inputs. Many computational approaches recently developed to derive drug binding kinetics^12–15^ exist and they can be used to estimate those parameters. In particular, the SSB toolkit has an implementation of the tRAMD tool^16^. This tool uses random accelerated molecular dynamics (RAMD) simulations to calculate the residence time of a ligand, from which the *k*_*off*_ parameter can be obtained^16^. If *K*_*d*_ is supplied, the concentration of activated receptors is calculated according to the occupancy theory, while for given *k*_*on*_ or *k*_*off*_ the concentration of activated receptors is calculated directly from the ligand-receptors binding equation’s reaction.

Likewise, the simulation and pathway parameters must be set up. The SSB toolkit allows the user to adjust each parameter of the pathway: initial species concentrations and kinetic parameters, or to stick with the default parameter values.

To assist the computational biology community to understand all the potentialities of the SSB toolkit we have prepared a series of *Jupyter* notebooks tutorials^17^ that can be easily run in cloud computing environments directly from the browser. All the toolkit source-code and the *Jupyter* notebook tutorials are open-source and available through *github*.*com* (https://github.com/rribeiro-sci/SSBtoolkit.git).

## Application Case: simulation of the signal-transducing pathway of the Oxytocin receptor (OXTR)

We have applied the SSB toolkit to model the signaling pathway of Oxytocin receptor (OXTR) in order to study the impact of a disease-variant of the receptor on its subcellular Ca^2+^ signaling dynamics^*5*^. We have implemented the OXTR-G_q_ signaling cascade by integrating two existing mathematical models, and by tuning the parameters of the model that were recognized as altered during disease conditions we were able to predict the functional impact caused by their associated variants^*5*^. Therefore, scaling one of the parameters of the model that describes the binding of the G_q_-protein to the receptor, which implicitly depends on the receptor activation, we were able to reproduce the shape of the experimental Ca^2+^ concentration curves. The integration of structural features within our model, allowed us to reproduce and give a rationale on the molecular determinants underlying the increase in the intracellular calcium concentration in presence of the variant. The results of the multiscale model prompted us to propose that the A218T variant is involved in a change in the activation properties of the OXTR with an impact on the downstream events. The SSB model of the OXTR, as well as other application cases, i.e. simulation of the agonist and competitive antagonists are described in detail in the Jupyter notebooks tutorials (https://github.com/rribeiro-sci/SSBtoolkit.git).

### Application Case: simulation of the dose-response curves of known antagonists and antagonists of Adenosine 2A receptor

We have applied the SSBtoolkit routines to model the cAMP concentration of the adenosine receptor (A2AR) upon binding of adenosine - the endogenous ligand - and the two more common agonists for A2 receptors used in functional studies: NECA and NGI to evaluate if we could reproduce experimental potency patterns. The experimental affinity values were obtained directly from *pubchem*.*org*^18^. To initiate the SB simulations, we have used *K*_*d*_ values of, obtained with a 3D-Convolution Neural Network Deep Learning (**Kdeep**)^19^ embedded in the *PlayMolecule*.*org* web server, from ligand-receptor (PDB ID: 2YDV) docked complexes performed with Autodock Vina^20^. We used a receptor concentration of 2 µM, a range of ligand concentration between 10-3 µM and 103 µM, and an integral time step of 1000. The results obtained from our protocol (Fig. S1) predicted that the NGI is the most potent agonist, followed by the NECA, being the Adenosine the less potent ligand, which agrees with the experimental data, Fig. S1. The same protocol was also applied to a set of antagonists of the A2 receptors, where we could observe that the predicted inhibition pattern also followed the experimental affinity pattern, Fig S2.

### Conclusions and future directions

Here we present an open-source toolkit designed to exploit the synergies between molecular-level ligand-GPCR interactions and higher-level systemic and physiologic mechanisms. What we are proposing not only might constitute a computational approach to systematically assess pharmacodynamic parameters during virtual screening campaigns against GRCRs, but also open the possibility to consistently model experimental data in order to give rationales on the molecular structure/function relationships of GPCRs, as well as on the impact of their disease associated variants along the signaling pathway.

Therefore, the insights obtained from our systems biology methods combined with a molecular level modeling of disease associated variants may support strategies for drug discovery and biomedical projects, paving the way to the acclaimed paradigm of *structural systems pharmacology*.

Even if the SSB toolkit was developed having in mind ligand-GPCRs systems, it can be expanded to other pharmacological systems. By implementing ODEs models of other signaling pathways^4^ it will be possible to cover a larger pharmacology landscape, and then, enhance our understanding in drugs mode of actions and increase the predictive power of pharmacokinetics/dynamics models.

## Methods

Since the SSB toolkit is an ongoing project, we strongly recommend users to read the full detailed documentation in the source-code repository on GitHub: https://github.com/rribeiro-sci/SSBtoolkit.git. Nonetheless, in the following lines a brief description on the methodology used to obtain drug-response curves for agonists and antagonists is provided.

### Obtaining dose-response curves of ligand-target interactions from simulation of mathematical models of signaling pathways

One of the key elements of the simulation of a mechanistic model of a signaling pathway is the calculation of the concentration of activated receptors under a predefined concentration of receptor and ligand. Typically, it is assumed that the response of a drug is proportional to the fraction of activated receptors. However, this assumption is not valid for the so-called “spare receptors” like GPCRs. That is to say that the maximum response of a GPCR can be achieved with less than 100% of occupancy^21^. As this implies, we can’t estimate the EC_50_/IC_50_ of ligands of GPCRs without including the signaling pathway associated with these receptors.

And there are two methods to calculate it. Having experimental or computational affinities (*K*_*d*_) the concentration of activated receptors can be calculated according to the occupancy theory that states that the concentration of activated receptors equals to the fraction of occupied receptors at equilibrium. On the other hand, the concentration of activated receptors can also be calculated directly using the ligand-target reaction equation if its kinetic parameters (*k*_*on*_ and *k*_*off*_) are provided.

With the *K*_*d*_, *k*_*on*_ or *k*_*off*_ parameters simulations of the mathematical models of the signaling pathway can be performed, by integrating all ODEs over a range of time. The variation of the concentration of the molecular species that make part of the model are, then, calculated from the time evolution of the ODEs. The response of a signaling pathway is, naturally, represented by the increase or decrease of one of the species described by the model. Therefore, for each signaling pathway we defined, by default, a reference species. While cAMP is chosen, by default, as reference species for the G_s_ and G_i/o_ pathway, for the G_q/11_ pathway we chose IP_3_ as default molecular species. However, the user has the freedom to select any of the signaling species included in the pathways for the data collection.

Finally, to obtain a dose-response curve from systems biology simulations, individual simulations of the model according to range of ligand concentrations must be performed first, as it is described in the main text.

However, this method is just valid if the ligands act as agonists. If the ligands act as competitive antagonists, the concentration of activated receptors will depend on the concentration and dissociation values of both the agonist and antagonist. By definition, an antagonist is a drug that inhibits the action of an agonist, having no effect in the absence of the agonist^21^. To address this situation, it is necessary to obtain at first place the drug-receptor binding curve of an antagonist in the presence of an agonist, like it is done in radio-ligand experimental binding assays. In such assays, a competitive ligand called “displacer” competes with a ligand with radioactivity previously added to the system. Therefore, we compute the following equation in function of a range of antagonist concentration with a saturation concentration value for the agonist:

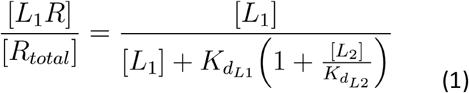

where L_1_R/R_total_ is the fraction of occupied receptors by agonist, L_1_ and K_dL1_ are the concentration and the dissociation constant of the agonist (‘labeled ligand’), L_2_ and K_dL2_ are the concentration and dissociation constant of the antagonist (‘displacer’).

To calculate the concentration of agonist that saturates the receptor, we must calculate its submaximal concentration, i.e., the concentration of agonist for which the fraction of occupied receptor reaches the maximum plateau on the agonist-receptor binding curve. Therefore, before obtaining a drug-receptor binding curve of the agonist in the presence of the antagonist, a drug-receptor binding curve of the agonist without the antagonist must be obtained applying a simplified version of the previous equation:

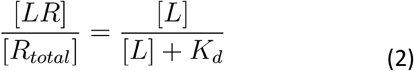

The submaximal concentration value for the agonist can then be deduced from the binding curve of the agonists using the methods proposed by *Sebaugh et al*.^22^, in which the maximum plateau on the agonist-receptor binding curve is defined as the upper bend point of the linear portion of the sigmoid curve that is mathematically obtained by calculating the maximum value of the derivative function of the logistic function.

## Availability

Source-code: https://github.com/rribeiro-sci/SSBtoolkit.git

Documentation:

Tutorials: https://colab.research.google.com/github/rribeiro-sci/SSBtoolkit/blob/main/SSBtoolkit-Tutorial1.ipynb; https://colab.research.google.com/github/rribeiro-sci/SSBtoolkit/blob/main/SSBtoolkit-Tutorial2.ipynb; https://colab.research.google.com/github/rribeiro-sci/SSBtoolkit/blob/main/SSBtoolkit-Tutorial3A.ipynb; https://colab.research.google.com/github/rribeiro-sci/SSBtoolkit/blob/main/SSBtoolkit-Tutorial3B-tauRAMD.ipynb https://colab.research.google.com/github/rribeiro-sci/SSBtoolkit/blob/main/SSBtoolkit-Tutorial4-OXTR.ipynb

