## Supplementary figures for "SSB toolkit: from molecular structure to subcellular signaling pathways"

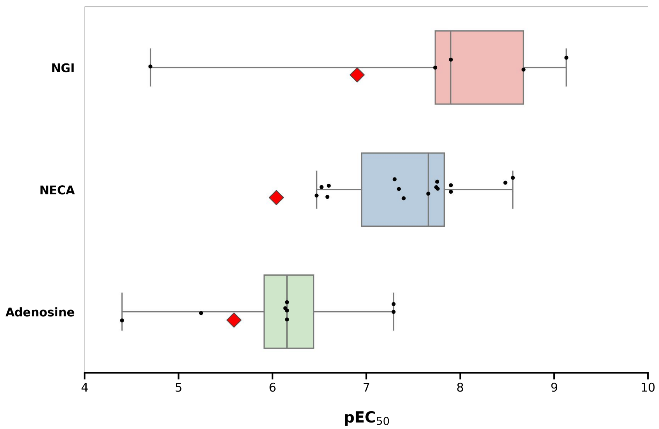


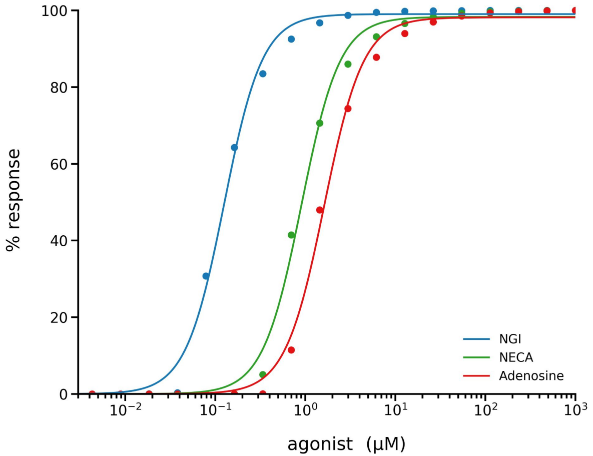


Fig S2 - (A) Statistical distribution of the experimental values of EC50 relative to the interaction between the adenosine, NECA and NGI with the A2A receptor, respectively. Data taken from PubChem.org. The red diamond marks represent the predicted values obtained from the SSB toolkit. (B and C) Dose-response curves of agonists (Adenosine, NECA and NGI) of A2A receptor and predicted EC50 values obtained with the SSB toolkit.

| Ligand | Predicted $EC_{50}$ (μM) | Predicted $pEC_{50}$ | Experimental* $pEC_{50}$ |
|---|---|---|---|
| Adenosine | 1.62 | 5.79 | 6.15 |
| NECA | 0.9 | 6.04 | 7.66 |
| NGI | 0.13 | 6.90 | 7.90 |

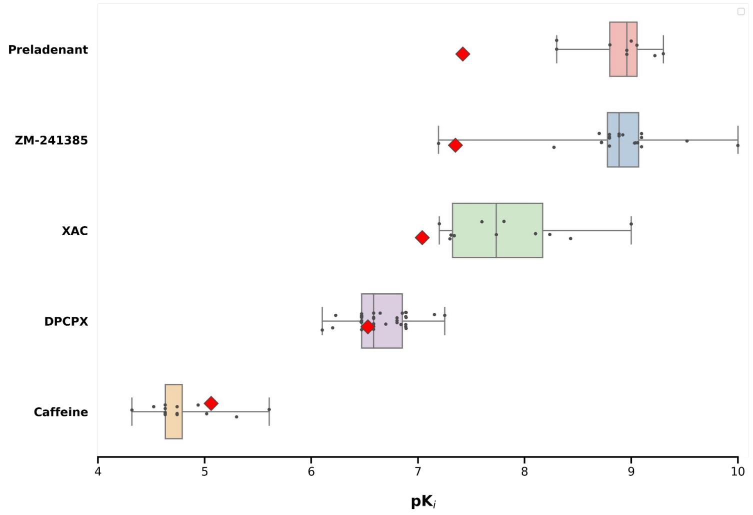


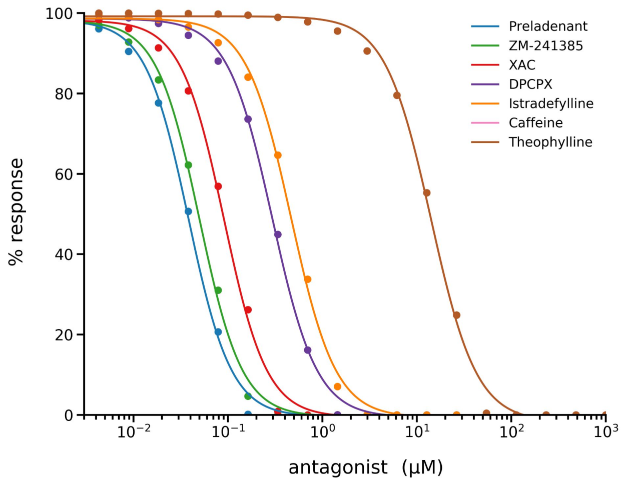


Fig S2 - (A) Statistical distribution of the experimental values of pKi relative to the interaction between antagonist with the A2A receptor. Data taken from PubChem.org. The red diamond marks represent the predicted values obtained from the SSB toolkit. (B and C) Dose-response curves of the antagonists and predicted IC50 values obtained with the SSB toolkit.

| Ligand | Predicted $IC_{50}$ (μM) | Predicted $pIC_{50}$ | Experimental* *pKi* |
|---|---|---|---|
| Caffeine | 8.76 | 5.06 | 4.63 |
| DPCPX | 0.29 | 6.53 | 6.58 |
| XAC | 0.09 | 7.04 | 7.74 |
| ZM-241385 | 0.05 | 7.30 | 8.89 |
| Preladenant | 0.04 | 7.42 | 8.96 |
